# Conjunctival microbiome-host responses are associated with impaired epithelial cell health in both early and late stages of trachoma

**DOI:** 10.1101/670711

**Authors:** Harry Pickering, Christine D Palmer, Joanna Houghton, Pateh Makalo, Hassan Joof, Tamsyn Derrick, Adriana Goncalves, David CW Mabey, Robin L Bailey, Matthew J Burton, Chrissy h Roberts, Sarah E Burr, Martin J Holland

**Author notes:** Corresponding author: Harry Pickering. Contributed equally (Harry Pickering, Christine D Palmer, Joanna Houghton).

## Abstract

**Background:** Trachoma, a neglected tropical disease, is the leading infectious cause of blindness and visual impairment worldwide. Host responses to ocular chlamydial infection resulting in chronic inflammation and expansion of non-chlamydial bacteria are hypothesised risk factors for development of active trachoma and conjunctival scarring

**Methods:** Ocular swabs from trachoma endemic populations in The Gambia were selected from archived samples for 16S sequencing and host conjunctival gene expression. We recruited children with active trachoma and adults with conjunctival scarring, alongside corresponding matched controls.

**Findings:** In children, active trachoma was not associated with significant changes in the ocular microbiome. *Haemophilus* enrichment was associated with antimicrobial responses but not linked to active trachoma. Adults with scarring trachoma had a reduced ocular bacterial diversity compared to controls, with increased relative abundance of *Corynebacterium*. Increased abundance of *Corynebacterium* in scarring disease was associated with innate immune responses to the microbiota, dominated by altered mucin expression and increased matrix adhesion.

**Interpretation:** In the absence of current *C. trachomatis* infection, changes in the ocular microbiome associate with antimicrobial and inflammatory responses that impair epithelial cell health. In scarring trachoma, expansion of ‘non-pathogenic’ bacteria such as *Corynebacterium* and innate responses are coincident, warranting further investigation of this relationship. Comparisons between active and scarring trachoma supported the relative absence of type-1 interferon responses in scarring, whilst highlighting a common suppression of re-epithelialisation with altered epithelial and bacterial adhesion, likely contributing to development of scarring pathology.

## Introduction

Ocular *Chlamydia trachomatis* (*Ct*) infection causes trachoma, the leading infectious cause of blindness worldwide. The pathophysiology of trachoma is complex and multifactorial.^1, 2^ The factors involved in the inflammatory responses to repeated *Ct* infection that lead to conjunctival scarring, trichiasis, corneal opacity and blindness remain poorly understood. In addition to *Ct* infection, other factors, including the type and quality of the conjunctival host immune responses,^3-6^ host genetic background,^7^ infections with other ocular pathogens and changes in overall bacterial community composition^8-11^ have each been linked to the different stages of trachomatous disease. Thus far there have been a limited number of studies that have investigated the interaction between the non-chlamydial ocular microbiota and conjunctival immune response in trachoma.^12, 13^

Culture-dependent methods have been used extensively to study the ocular surface microbiome, the first descriptions dating back to 1930.^14^ Initial reports generally considered between 20 and 80% of normal healthy eyes to be sterile. The ocular surface is still generally considered to harbour a paucibacillary community, although bacteria have been isolated from higher proportions of healthy conjunctivae by using more intensive modern culture techniques.^15^ Efforts to define the normal ocular flora in African populations were initially conducted in rural Sierra Leone. The authors found several microbial species, including *Staphylococcus* spp., *Pseudomonas* spp., other Gram-negative populations and fungal species.^16^ Subsequent studies utilizing approaches to sequence prokaryotic 16S ribosomal RNA genes (16S rRNA) for identification of ocular bacterial communities confirmed the presence of *Staphylococcus* and *Pseudomonas* spp., amongst others, and further developed our understanding of the ocular microbiome beyond those species detectable by *in vitro* culture. Specifically, studies of the healthy human conjunctival microbiome have consistently identified *Pseudomonas* spp., *Propionibacterium* spp., *Acinetobacter* spp., *Corynebacterium* spp., *Staphylococci, Micrococcus* spp. and *Streptococci*.^17-21^

Congruent with observations in other anatomical sites, several studies proposed a link between the ocular microbiota, ocular health and susceptibility to infections. Using culture-dependent techniques, bacteria were more frequently identified and at higher abundance in samples from bacterial conjunctivitis patients compared to healthy controls. However, cultured species predominantly overlapped with those identified by others as present in healthy conjunctival samples.^22^ Two recent studies in murine models of ocular infection have demonstrated an important role for the ocular microbiota in bolstering local immune responses and increasing resistance to infectious challenge. Firstly, *Pseudomonas aeruginosa*–induced keratitis resistant mice became susceptible in the absence of ocular microbiota. The protection afforded was mediated by a microbiota-induced IL-1β-dependent mechanism.^23^ Secondly, a constituent of the ocular commensal flora, *Corynebacterium mastitidis*, was shown to mediate protection against ocular fungal (*Candida albicans)* and bacterial *(P. aeruginosa)* challenge infection in mice. In this case, *C. mastitidis* was found to elicit a local IL-17 response that was central to neutrophil recruitment and release of antimicrobials into the tears, leading to increased resistance.^24^ In addition to stimulating local immune responses, *Corynebacterium* spp, which are consistently found as a major constituent of the ocular microbiome, may also protect their ecological niche against specific pathogens such as *Streptococcus pneumoniae* (*Sp*) via release of antibacterial free fatty acids that inhibit their growth.^25^

In typical cases of active trachoma with proven *Ct* infection, such as reported in historical studies in The Gambia, the conjunctival response to *Ct* infection is characterized by epithelial cell reorganisation, immune cell infiltration and secretion of anti-microbial peptides.^3^ Similarly, trachomatous inflammation follicular (TF) and trachomatous scarring (TS) were also associated with expression of innate pro-inflammatory markers in Ethiopian and Tanzanian populations.^5, 6, 8, 13^ However, clinical signs of trachoma are often prevalent in the relative absence of *Ct* infection.^8, 26-28^ Additionally, longitudinal cohort studies in Tanzania and Ethiopia in adults with progressive conjunctival scarring found that concurrent *Ct* infection was virtually absent.^11^ In these studies, non-chlamydial bacteria are often prevalent and associate with trachomatous disease. These non-chlamydial bacteria include pathogens, such as *Sp* and *Haemophilus influenzae* (*Hi*), and commensals, such as *Corynebacterium spp*.^8, 9, 11, 26^ Non-chlamydial infections have also been associated with immunofibrogenic immune responses thought to drive scarring trachoma.^13^ Overall, studies from different trachoma endemic populations suggest that non-chlamydial bacterial species are significant factors in trachoma pathogenesis.^8, 11, 13, 26^

Using culture independent methods, we have previously shown differences in conjunctival microbiome diversity between individuals with trachomatous scarring and controls, with elevated abundance of *Corynebacterium* in adults with scarring and trichiasis.^10^ Here, we investigate the relationship between the conjunctival microbiome and host conjunctival-associated lymphoid tissue responses, additionally testing the influence of host genotype on the ocular microbiome in different clinical stages of trachoma in Gambians. Our data demonstrate significant associations between ocular microbiota the host immune-response linked to specific bacteria and trachomatous disease.

## Materials and Methods

### Ethics statement

The study was conducted in accordance with the Declaration of Helsinki. Permission for collection of samples and genotyping was granted by the relevant local and national ethics committees of the London School of Hygiene and Tropical Medicine, The Gambian Government/Medical Research Council Unit and The Gambia Joint Ethics Committee. Written, informed consent prior to a participant’s enrolment was obtained from all adult participants and from a parent or a guardian for participants under 18 years of age.

### Study populations

Samples (n=361) were selected based on case-control status (trachoma clinical signs) from a larger archive of ocular swabs collected from individuals in communities across The Gambia, West Africa between 2009 and 2011. Conjunctival microbiome data for a subset of samples (n=220) has been published previously.^10^ Cases of active or scarring trachoma were identified from screening records, community ophthalmic nurse referral and opportunistic rapid screening. Control individuals with normal conjunctivae were selected by matching for age, sex, ethnicity and location. Samples were classified as collected during the Gambian dry season (December – April) or wet season (July-October). No samples were collected in May, June or November of any year. Subject demographics are shown in Table 1.

**Table 1.** Demographic characteristics of participants

| Phenotype | Children (< 16 years of age) |  |  | Adults (≥ 16 years of age) |  |  |
| --- | --- | --- | --- | --- | --- | --- |
|  | Healthy<br>n = 36<br>(42%) | Active<br>trachoma n<br>= 49 (58%) | Adjusted<br>p-value | Healthy<br>n = 121<br>(43%) | Scarring<br>trachoma n =<br>158 (57%) | Adjusted<br>p-value |
| **Male (%)** | 22 (61) | 30 (61) | 1.000 | 35 (29) | 38 (22) | 0.435 |
| **Median age (range)** | 6 (1-14) | 5 (1-13) | 0.275 | 53 (16-87) | 55 (16-84) | 0.119 |
| **Sample collection [n (%)]** |  |  | 1.000 |  |  | 1.000 |
| Dry season | 7 (19) | 9 (18) |  | 91 (75) | 118 (75) |  |
| Wet season | 29 (81) | 40 (82) |  | 30 (25) | 40 (25) |  |
| **Ethnicity [n(%)]** |  |  | 0.445 |  |  | 0.938 |
| Jola | 6 (17) | 6 (12) |  | 37 (31) | 48 (30) |  |
| Mandinka | 14 (39) | 17 (35) |  | 56 (46) | 76 (48) |  |
| Wolof | 11 (31) | 12 (24) |  | 10 (8) | 10 (6) |  |
| Other | 5 (14) | 14 (29) |  | 18 (15) | 24 (15) |  |
| **Administrative district [n (%)]** |  |  | 0.102 |  |  | 0.922 |
| Basse | 0 (0) | 0 (0) |  | 15 (12) | 21 (9) |  |
| Brikama | 11 (31) | 15 (31) |  | 58 (48) | 79 (37) |  |
| Janjanbureh | 1 (3) | 0 (0) |  | 2 (2) | 4 (1) |  |
| Kanifing | 0 (0) | 0 (0) |  | 1 (1) | 1 (1) |  |
| Kerewan | 0 (0) | 3 (6) |  | 9 (7) | 12 (6) |  |
| Kuntaut | 3 (8) | 0 (0) |  | 7 (6) | 6 (4) |  |
| Mansa Konko | 21 (58) | 31 (63) |  | 19 (16) | 35 (22) |  |
| Unknown | 0 (0) | 0 (0) |  | 10 (8) | 0 (0) |  |

### Ocular swab collection

Swab samples were taken from the everted left and right tarsal conjunctiva of each study participant using standard methodology.^29, 30^ A separate swab was used for each eye and each swab was horizontally passed across the tarsal conjunctiva three times, rotating the swab by 1/3 with each pass. Individual swabs were immediately placed into sterile tubes filled with 250μl RNAlater® (Ambion) and stored in a cool-box filled with ice packs in the field, then transferred to -80°C storage in the laboratory.

### Photographs and clinical scoring

Subjects were examined for clinical signs of trachoma in the field. High resolution digital photographs were taken of each conjunctival surface at the time of sample collection and an FPC score (1981 WHO Trachoma Grading System; FPC – follicles, papillae, cicatricae^31^) assigned to each sample by two experienced trachoma graders, as described previously.^10^ For analyses in this study, the presence of follicles was defined as an F score > 0. Presence of papillae (indicative of inflammation) was graded as a P score > 0. Conjunctival scarring was defined as a C score > 0. Participants with normal, healthy conjunctivae, as defined by a score of F0|P0|C0, served as controls.

### Genomic DNA and RNA extraction from ocular swabs

Ocular swabs were removed from RNALater® and placed into 400 µl Norgen lysis buffer (Norgen Biotek), vortexed for 1 minute and spun down for 30 seconds at 13,000 rpm. Lysates were transferred into a new RNAse/DNAse free tubes. Extraction of nucleic acid material was performed using the Norgen total RNA/DNA purification kit according to the manufacturer’s instructions. β-mercaptoethanol was added to lysis buffers to inhibit RNAses. RNA and gDNA were stored at -80° C until further processing.

### cDNA synthesis

Generation of cDNA from RNA was performed using the SuperScript® VILO cDNA Synthesis Kit and Master Mix (Invitrogen) according to the manufacturer’s instructions. Total RNA-derived cDNA (which includes cDNA derived from miRNA) was generated using the miScript II RT Kit (Qiagen) using the hiFlex buffer according to the manufacturer’s instructions. cDNA was used immediately or stored at -20° C until further processing.

### Taqman® Low Density Array (TLDA)

Analysis of ocular swab cDNA was performed using TLDA Microfluidic Cards (Applied Biosystems) according to the manufacturer’s instructions. Custom array cards were used to interrogate expression profiles of immune transcripts (Supplementary Table 1). Arrays were run on an ABI PRISM® 7900HT thermal cycler (Applied Biosystems) using SDS software. Cycling conditions were as follows: 2 min at 50° C | 10 min at 94.5° C | 40 cycles (30 sec at 97° C | 1 min at 59.7° C). Data were collected at 97° C and 59.7° C. Cycle threshold (C_T_) was set to a standard mid-exponential phase amplification point. Delta C_T_ values relative to housekeeping genes (*GAPDH* and *HPRT1*) were calculated using R (R: A language and environment for statistical computing. R Foundation for Statistical Computing, Vienna, Austria; https://www.R-project.org). Genes and individuals with ≥ 10% missing data were excluded. Fold-change was calculated using the delta-delta CT method. Modules of co-expressed genes were identified using the Weighted Correlation Network Analysis (WGCNA) package and putative functions were assigned manually based on known functions of genes in each module.^32, 33^ All genes in each module were included in a principal component analysis (PCA), the first component of which was used as an expression score per individual for each module.

### miRNA qPCR

qPCR was carried out using miScript Primer Assays and the miScript SYBR Green PCR kit according to the manufacturer’s instructions (Qiagen) and data were acquired on an ABI PRISM® 7900HT thermal cycler using SDS software. Primer assays used were: Hs_RNU6-2_11 (MS00033740), Hs_miR-1285_2 (MS00031367), Hs_miR-147b_1 (MS00008729), Hs_miR-155_2 (MS00031486) and Hs_miR-184_1 (MS00003640). Cycling conditions were as follows: 15 min at 95° C | 40 cycles (15 sec at 94° C | 30 sec at 55° C | 30 sec at 70° C). Data were collected at 94° C and 70° C. C_T_ was determined automatically, or adjusted manually to cross amplification curves in the mid-exponential phase where not placed correctly by the software. Delta C_T_ values relative to housekeeping genes (Hs_RNU6-2_11) were calculated using R.

### Quantification of bacterial and human DNA by Droplet Digital™PCR

Detection and quantitation of *Chlamydia trachomatis* (*Ct*) plasmid and human DNA in ocular swab samples was performed using the Bio-rad QX100™droplet digital PCR (ddPCR) platform. Primers and protocols for the specific amplification of *Homo sapiens* ribonuclease P/MRP 30kDa subunit (*RPP30*), *Ct* plasmid, *Haemophilus influenzae* and *Streptococcus pneumoniae* were all described previously.^27, 34-36^

### Prokaryotic 16S ribosomal RNA gene (16S rRNA) sequencing

Ocular bacterial community composition was determined previously by 454 pyrosequencing.^10^ Additional samples were analysed by 16S rRNA sequencing using the MiSeq Next Generation Sequencing Platform (Illumina). Amplicon PCR was performed using the Phusion High-Fidelity PCR Master Mix (New England Biolabs) and previously described barcoded primers^37^ to amplify the V1-V3 region of bacterial 16S rDNA. Cycling conditions were as follows: 30 sec at 98° C | 31 cycles (10 sec at 98° C | 30 sec at 62° C | 15 sec at 72° C) | 7 min at 72° C | hold at 4° C. PCR products (∼600 bp) were confirmed in a subset of samples using agarose gel electrophoresis. Purification of PCR products was performed using the Agencourt AMPure XP system (Beckman Coulter) and DNA quantified using the Qubit® 2.0 Fluorometer according to the manufacturer’s instructions (ThermoFisher Scientific). Samples were pooled into a DNA library, which was denatured and run on the MiSeq sequencer at a final concentration of 5 pM alongside a 5 pM PhiX control (Illumina). Raw reads generated by MiSeq were error-corrected and filtered using DADA2 through QIIME2 (https://qiime2.org).^38^ Filtered reads were clustered *de novo* into Operational Taxonomic Units (OTUs) at 97% sequence similarity. OTUs were then assigned taxonomy using a Naive Bayes classifier trained on the SILVA 16S database. Both processes were performed with QIIME2. Manual filtering of classified OTUs was performed using R as described below. OTUs were retained if they had been classified as bacteria, had a genus-level classification and constituted >0.005% of the total number of reads.^39^ Samples with <1000 reads were excluded. Final read counts were rarefied to 1000 reads per sample using the R package vegan. 16S rRNA sequencing data (genus counts) generated by Roche-454 (n=220) published previously^10^ were included in the analysis. Read counts in the combined datasets were converted to relative abundance of each phyla or genera per individual for all analyses. Univariate analyses included genera with an abundance >1%.

### KLRC2 genotyping

*KLRC2* genotypes were determined by touchdown PCR using the Phusion High Fidelity PCR kit (New England Biolabs) using previously described methods^40, 41^ and primers.^42^ Touchdown PCR was carried out as previously described.^41, 42^ Cycling conditions were as follows: 3 min at 95° C | 10 cycles (30 sec at 94° C | 30 sec from 65° C to 55° C reducing by 1° C per cycle) | 30 sec at 72° C | 26 cycles (30 sec at 94° C | 30 sec at 55° C | 30 sec at 72° C). PCR products were separated and identified using agarose gel electrophoresis.

### KIR copy number variation (CNV) assay

Determination of *KIR2DL2* and *KIR2DL3* alleles was performed by ddPCR (BioRad) on buccal brush extracted DNA.^43^ Each allele was tested for separately in parallel with the human target *RPP30*. Primers and probes were master mixed to final concentration of 3 µM each of primer and 1 µM of probe except in the case of *KIR2DL2* (0.5 µM). Primer sequences were as previously described.^7^ Samples were digested with 1 unit of BamHI-HF enzyme (NEB) prior to running the PCR. PCR reactions contained 4µl of digested sample, 2 µl of primer probe master mix, 4 µl molecular grade water and 10 µl of 2x ddPCR master mix (BioRad). Cycling conditions were as follows: 10 min 95°C | 43 cycles (15 sec 95° C | 60 sec 60° C) | 12 min 98° C.

### HLA-C typing

Extracted DNA from ocular swabs was used for HLA-C1/C2 epityping by allelic discrimination on an ABI 7900HT. Primer and probes used in the reaction were as follows; Forward primer HLA-C-JS_C1C2F 5’-TATTGGGACCGGGAGACACA-3’, Reverse primer HLA-C-3C26-R 5’-GGAGGGGTCGTGACCTGCGC-3’, C1 probe BARI-C1 6FAM-CCGAGTGAGCCTGC-MGBNFQ, C2 probe BARI-C2 VIC-CCGAGTGAACCTGC-MGBNFQ.^44^ Each PCR reaction contained 3ul Taqman genotyping mastermix (Applied Biosystems), 1.7ul molecular grade water, 0.3ul primer and probe mix (JS C1C2F, HLA-3C26, Bari-C probes, all at 1uM) and 1µl template. Cycling conditions were as follows: 10 min at 95°C | 50 cycles (15 sec 95° C | 60 sec 60° C). Allele calls were made using SDS software v2.4.

### Corynebacterium rpoB amplification and sequencing

DNA samples that were positive for *Corynebacterium* from 16S sequencing were amplified using degenerate primers C2700 and C3130.^45^ Briefly, the PCR mix consisted of 1 x Ultra-red mix (PCR Biosystems), 400 nM of each primer and 2 µl of template DNA. Cycling conditions were as follows: 1 min at 95° C | 6 cycles (15 sec at 95° C | 15 sec at 68-62° C touchdown | 30 sec at 72° C) | 35 cycles (15 sec at 95° C | 15 sec at 62° C | 30 sec at 72° C) | 10 min at 72° C | hold at 4° C. PCR products were confirmed using agarose gel electrophoresis. Purification of PCR products was performed using the Agencourt AMPure XP system (Beckman Coulter) and DNA quantified using the Qubit® 2.0 Fluorometer according to the manufacturer’s instructions (ThermoFisher Scientific). ABI Prism terminator reactions were then performed as per manufacturer’s instructions, and consequently sequenced on the ABI 3700 capillary sequencer. Sequence data was analysed within R by inputting the .ab files and trimming the end N bases before quality filtering to Q10. Sequences were classified using blastn sequence identity.

### Bacterial diversity using Hill numbers

Global differences in the ocular microbiome were examined using Hill numbers, which take into account both richness and evenness (Equation 1).^46^ Where *S* is number of genera, *pi* is the proportion of genera per individual and *q* is the order of diversity. As the order of diversity (*q*) increases, greater weight is placed on the most abundant genera, reducing the Hill number. Within each value of *q*, higher values indicate increased diversity. Samples with few, dominant genera will have a lower Hill number, reflecting unevenness and reduced diversity. Samples with many, equally abundant genera will have a higher Hill number, reflecting increased diversity and evenness.

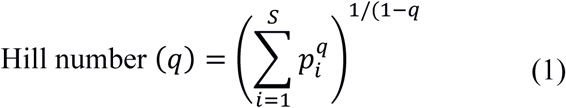

### Statistical analyses

R was used for statistical analyses and graphical visualizations. Analyses were performed using the linear model function to compute p-values unless otherwise stated. Analyses of gene expression (GE) were conducted using a linear regression of GE as the dependent variable and host disease phenotype as the independent variable, adjusted for age and gender. Analyses of the ocular microbiome were conducted using a linear regression of microbial diversity or relative abundance as the dependent variable and host disease phenotype as the independent variable, adjusted for age, gender and season. All analyses in adults were additionally adjusted for evidence of P-score > 0. P-values were adjusted using the false discovery rate (FDR) by the Benjamini-Hochberg procedure. Contingency analyses were performed by Chi-square test or Fisher’s exact test. Scaled PCA was performed in R using the *stats p*ackage. Hill numbers, diversity indices and nonmetric multidimensional scaling were calculated using the *vegan* package. P-values were considered significant at <0.05 and are denominated in figures as follows: * p < 0.05; ** p < 0.01; *** p < 0.001.

## Results

### Participants

Thirty-six children (< 16 years) with a normal, healthy conjunctiva (“N”; F0|P0) and 49 with active trachoma (“AT”; F > 0 +/- P > 0) were included in this study. Evidence of scarring was not significantly different between N (4/36 [11%]) and AT (9/49 [18%]). Of the adults studied, 121 (≥16 years) had a normal, healthy conjunctiva (“N”; F0|P0|C0) and 158 had scarring trachoma (“ST”; C > 0). Additionally, 76/158 (48%) adults with ST had P-score > 0, which was adjusted for in all analyses. No demographic variables were significantly associated with AT in children or with ST in adults, and unless otherwise stated, downstream analyses were only adjusted for age and gender.

### Gene expression patterns in active and scarring trachoma

Gene expression (GE) was characterised using TLDA Microfluidic Cards for targeted immune transcripts and qPCR for miRNA transcripts previously identified as associated with trachoma.^3, 5, 47, 48^ GE data were available from 78/85 children (N = 32, AT = 46) and 147/279 adults (N = 69, ST = 78) (Supplementary Table 2).

The expression of seven genes was significantly upregulated in AT (Figure 1A; *S100A7, DEFB4B, IL-17A, IL-23A, CD274, SRGN* and *CD53*). All seven genes were in the WCGNA defined GE module we termed “Active Trachoma Expression Module (ATEM) 2”, the contents of which are putatively involved in the innate response to microbiota (Supplementary Table 3). Combined expression level of ATEM2 was also significantly upregulated in active trachoma (adj.p = 0.002, coef = 2.267 [se = 0.704]). Four genes were significantly downregulated in AT (Figure 1A; *SPARCL1, TRAF6, MUC16* and *TNFSF15*). Similarly, all four genes were in the same GE module ATEM3, which is putatively involved in suppression of epithelial cell expansion and recovery (Supplementary Table 3).

**Figure 1.**
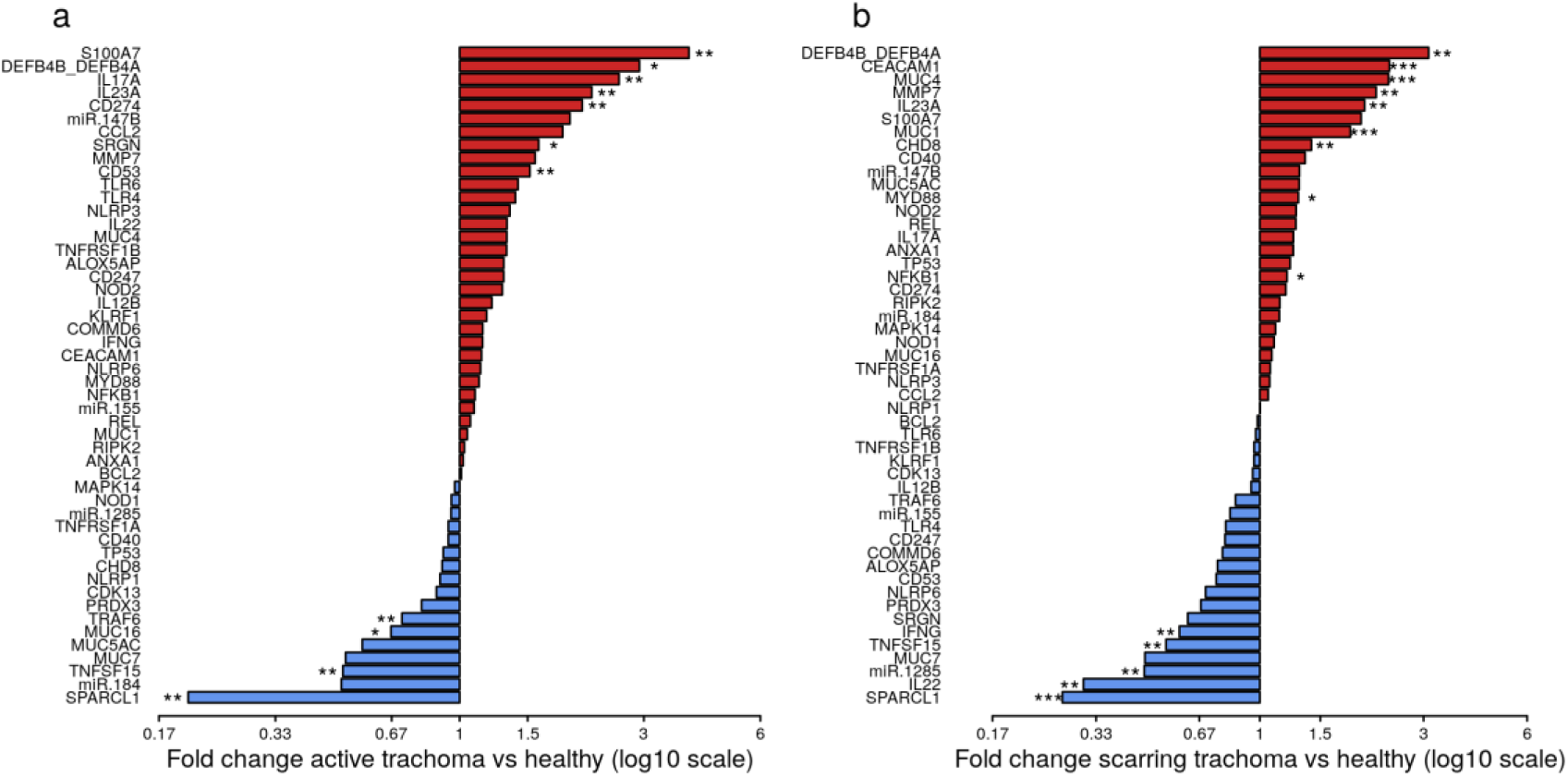
Fold-changes in conjunctival gene expression between cases and matched, healthy controls. Magnitude of fold-changes in conjunctival gene expression between children with active trachoma and healthy controls (a) and adults with scarring trachoma and healthy controls (b), shown by bars. Colours highlight increased (red) or decreased (blue) expression in cases. P-values were considered significant at <0.05 and are denominated as follows: * p<0.05; ** p<0.01; *** p<0.001.

In ST, the expression of nine genes was significantly upregulated (Figure 1B; *DEFB4B, CEACAM1, MUC4, MMP7, IL-23A, MUC1, CHD8, MYD88* and *NFKB1*). Six of nine genes were in GE module “Scarring Trachoma Expression Module (STEM) 4”, characteristically involved in responses to microbiota (Supplementary Table 4). Combined expression level of STEM4 was also significantly upregulated in ST (adj.p = 7.050 ×10^−7^, coef = 2.117 [se = 0.405]). Five genes were significantly downregulated in ST (Figure 1B; *SPARCL1, IL-22, miR-1285, TNFSF15* and *IFNy*). Three of four genes were in the same GE module STEM1, involved in epithelial health (Supplementary Table 4).

To identify shared differential gene expression in active and scarring trachoma, fold-changes from significantly differentially expressed genes in the previously described comparisons (Figure 1) were contrasted. This highlighted three groups of genes that were a) downregulated in AT and ST (Figure 2, blue hatched area), b) upregulated in AT and ST (Figure 2, red hatched area), and c) differentially regulated in AT and ST (Figure 2, green cross-hatched area). Co-downregulated genes (Figure 2; blue hatched area) are primarily involved in regulation of cellular expansion and migration. Of the differentially regulated genes (Figure 2, green cross-hatched area), those upregulated in AT and downregulated in ST are part of the host pro-inflammatory response. Co-upregulated genes (Figure 2; red hatched area) are mostly antimicrobial or pro-inflammatory. Within this latter set of co-upregulated genes, the magnitude of upregulation is higher in mucins and MMP7 in ST relative to AT. Conversely, the magnitude of upregulation is relatively higher in antimicrobials and pro-inflammatory IL-17A in AT.

**Figure 2.**
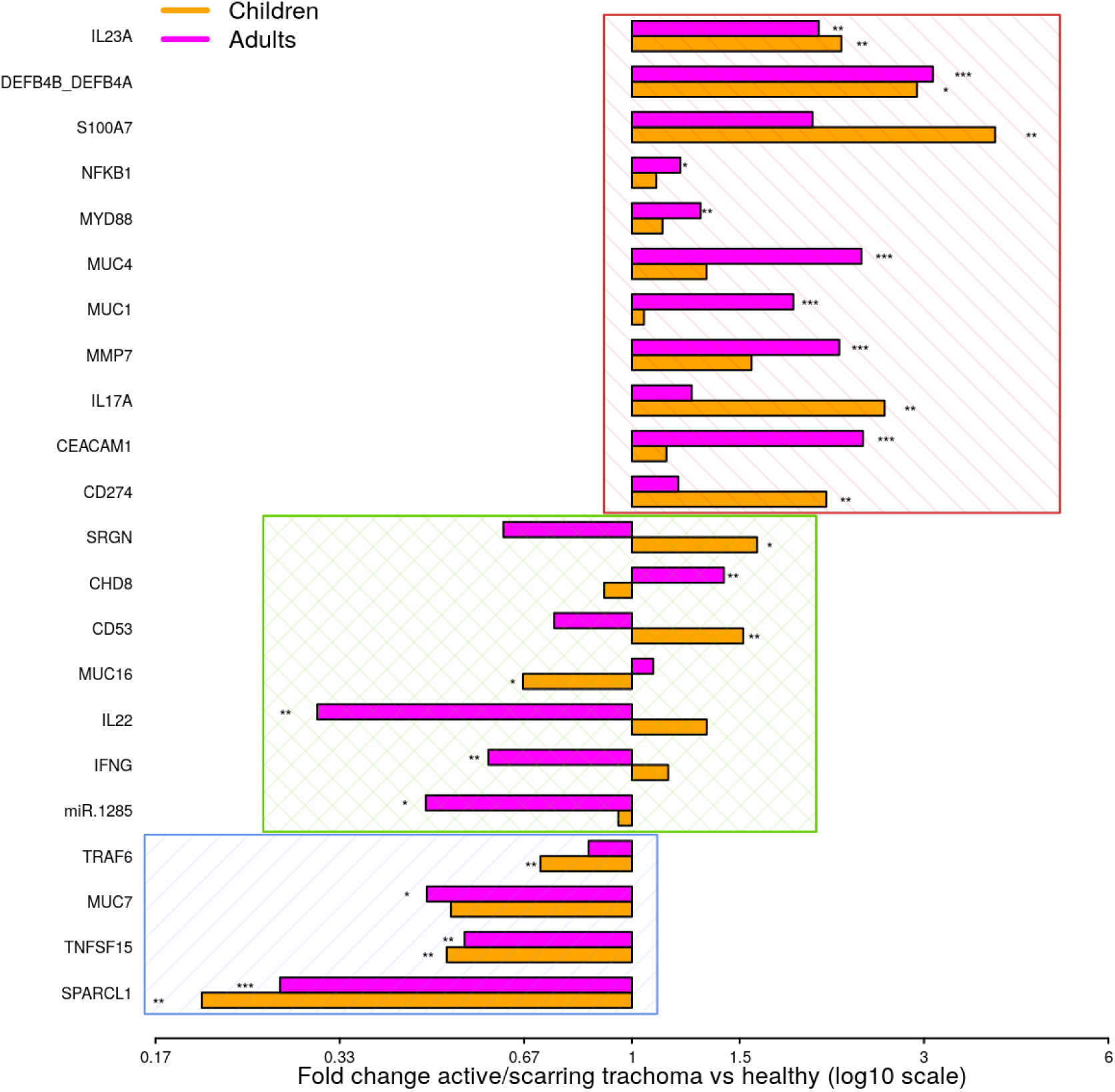
Comparison of fold-changes in conjunctival gene expression between active/scarring trachoma cases and healthy controls. Fold-changes in gene expression between children with active trachoma and healthy controls (orange bars) and adults with scarring trachoma and healthy controls (purple bars) are represented by bars with significance indicated as described below. Genes are sorted into three groups; downregulated in active and scarring trachoma (blue area), upregulated in active and scarring and trachoma (red area), and differentially regulated in active and scarring trachoma (green area). P-values were considered significant at <0.05 and are denominated as follows: * p<0.05; ** p<0.01; *** p<0.001.

### Ocular microbial changes in active and scarring trachoma

The ocular microbial community was characterised by sequencing of the V1-V3 region of the 16S gene, combining previously published data generated by 454 method^10^ and new data generated by MiSeq sequencing. Nonmetric multidimensional scaling of the compiled datasets showed no significant difference between samples sequenced on the two platforms (p=0.436). 16S data was available from 72/85 children (N = 31, AT =41) and 235/279 adults (N = 105, ST = 130) (Supplementary Table 5).

We identified 382 genera from 29 phyla. As found previously, the ocular microbiome community diversity decreased with age (p = 0.0007, coef = -0.007 [se = 0.002]). Phyla with an abundance >1% were identical between children and adults. *Actinobacteria* and *Firmicutes* accounted for the majority of the observed microbes (Figure 3). Eleven genera had relative abundance > 1% in both children and adults, ten genera were > 1% in children only and one genus was > 1% in adults only (Figure 4).

**Figure 3.**
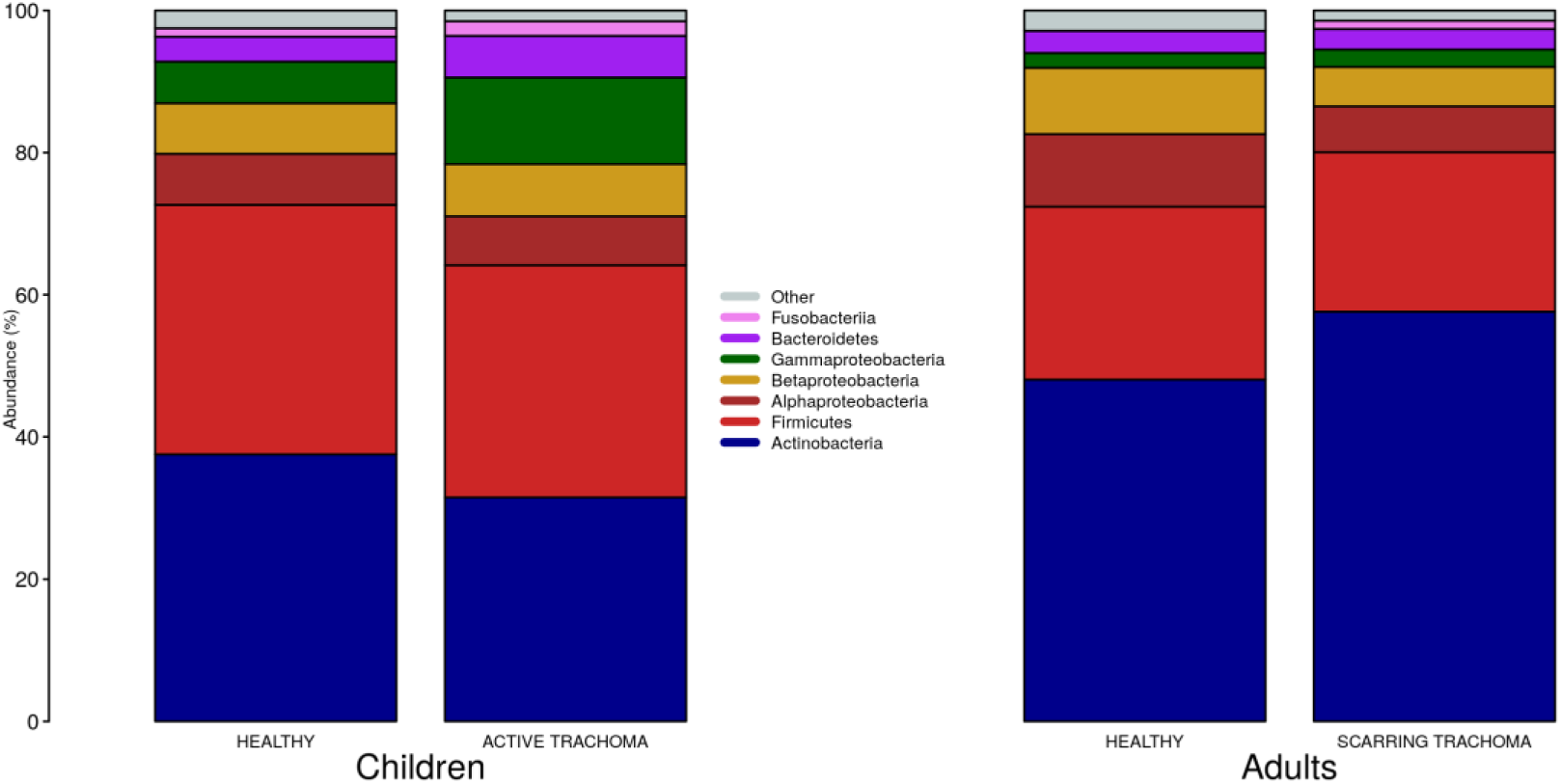
Relative abundance of major phyla in children and adults by case-control status. Phyla with relative abundance >1% in either children or adults are shown. Phyla with relative abundance ≤1% are grouped into ‘Other’.

**Figure 4.**
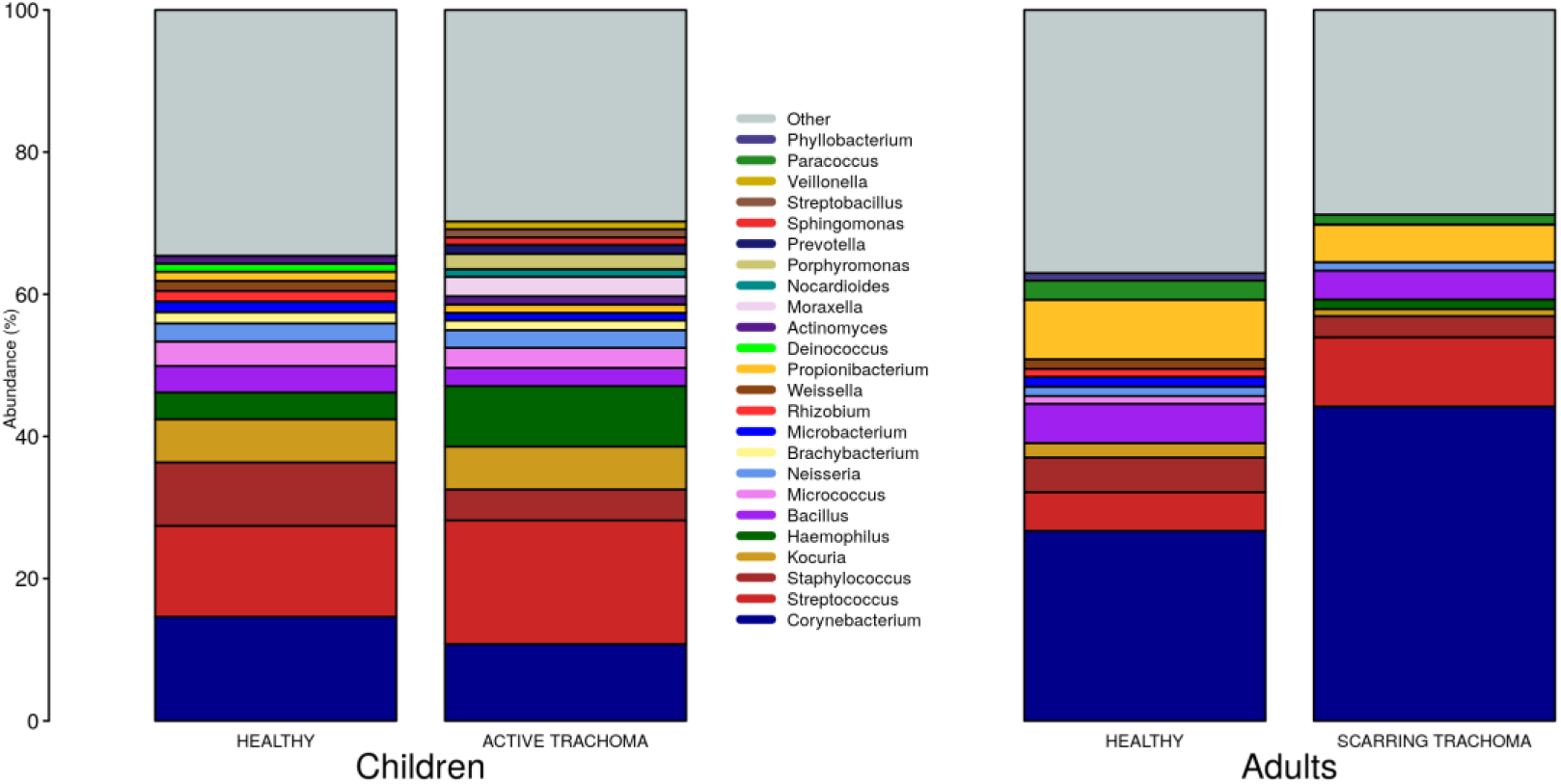
Relative abundance of major genera in children and adults by case-control status. Genera with relative abundance >1% in either children or adults are shown. Genera with relative abundance ≤1% are grouped into ‘Other’.

Global differences in the ocular microbiome were examined using Hill numbers, details are provided in the Methods. At order of diversity = 0 children with AT tended to have reduced diversity compared to healthy controls; Hill number did not significantly differ between AT cases and normal children when order of diversity was 0.5 or higher (Figure 5A). At order of diversity = 0, adults with ST were indistinguishable from normal, healthy adults (Figure 5B). From order of diversity = 0.5 upwards, adults with ST had significantly reduced diversity. The level of significance continued to increase with increasing order of diversity, suggesting dominance of a small number of genera.

**Figure 5.**
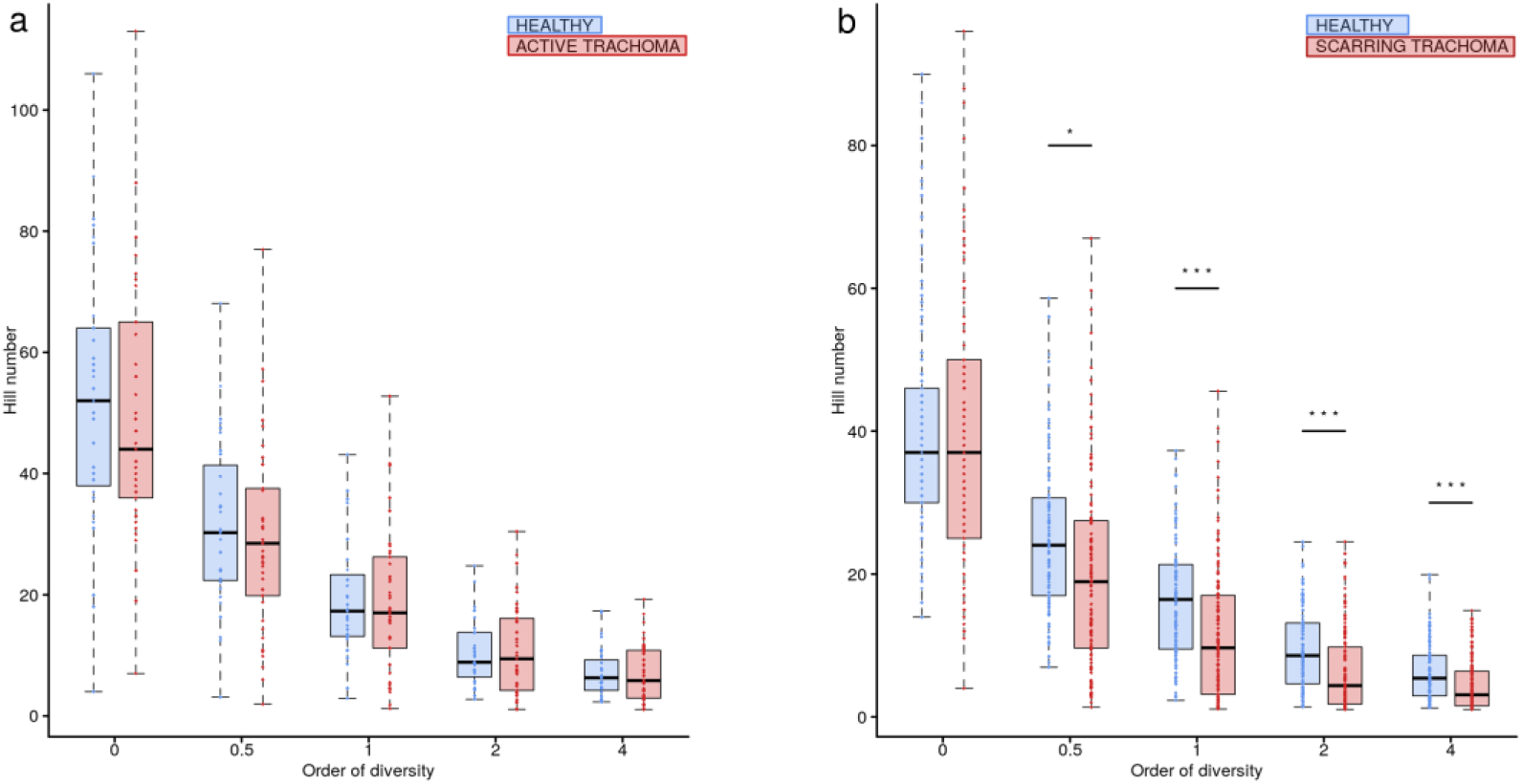
Ocular microbiome diversity in children and adults by case-control status. Hill number at corresponding order of diversity are shown for cases (red) and healthy controls (blue) in children (a) and adults (b). Boxes represent the interquartile range, with median indicated (blue line). Outer bars represent the range. P-values were considered significant at <0.05 and are denominated as follows: * p<0.05; ** p<0.01; *** p<0.001.

In univariate analyses there were no differentially abundant genera between children with AT and normal, healthy children. In adults with scarring trachoma, *Corynebacterium* abundance was significantly greater (adj.p = 0.0002, coef = 0.156 [se = 0.036]) and *Staphylococcus* abundance was significantly lower than in healthy controls (adj.p = 0.0011, coef = -0.022 [se = 0.006]) (Supplementary Figure 1). For both genera, prevalence (read count > 0) was equivalent between N and ST (*Corynebacterium*; N = 105/105 [100%] & ST = 129/130 [99%]. *Staphylococcus*; N = 90/105 [86%] & ST = 109/103 [84%]).

Individual direct tests for *Ct, Haemophilus influenzae (Hi*) and *S. pneumoniae (Sp)* were performed using ddPCR. *Hi* and *Sp* are non-chlamydial conjunctivitis-associated bacterial pathogens. In children, we identified seven cases of current *Ct* infection (7/82 [8.5%]). However, *Ct* infection was not associated with AT (p = 0.200 [N = 5/32, AT = 2/46]). No adults had detectable current *Ct* infection. The overall prevalence of *Hi* was 43/85 (51%) in children and 25/279 (9%) in adults. *Sp* prevalence was 28/85 (33%) in children and 12/279 (4%) in adults. In children, there were no differences in the proportion of AT cases and controls who were positive for *Hi* (p = 0.203 [N = 14/31, AT = 26/41]) or *Sp* (p = 0.333 [N = 8/31, AT = 18/41]). Using ddPCR as the reference, 16S sequencing had reasonable sensitivity but poor specificity (Table 2) for these bacteria. The high proportion of ddPCR-negative, 16S-positive samples suggests that additional members of the *Haemophilus* and *Streptococcus* genera were present on a significant number of conjunctivae. Correlation between ddPCR concentration and 16S relative abundance was also poor for both species (*Hi p* = 0.174, *Sp p* = 0.207). Abundance was higher however in PCR-positive samples, significantly so for *Sp* (*Hi* p = 0.081, *Sp* p = 0.002).

**Table 2.** Comparison of species-specific, ddPCR based test and 16S genus-level operational taxonomic unit (OTU) classification

|  | PCR negative (%) | PCR positive |
| --- | --- | --- |
| ***H. influenzae*** |  |  |
| 16S negative (%) | 57 (76) | 26 (46) |
| 16S positive (%) | 18 (24) | 31 (54) |
| Sensitivity (%) | 76 |  |
| Specificity (%) | 54 |  |
| ***S. pneumoniae*** |  |  |
| 16S negative (%) | 3 (3) | 0 (0) |
| 16S positive (%) | 97 (97) | 32 (100) |
| Sensitivity (%) | 100 |  |
| Specificity (%) | 3 |  |

Reads corresponding to the *Corynebacterium* OTU were detected in 332/364 (91.2%) samples. There was sufficient residual DNA for *Corynebacterium* specific PCR amplification and sequencing in 112/332 samples. Within the 112 samples that could be tested, there were 71 ST cases and 41 age matched controls, of which 18 samples (16 cases and two controls) passed Q10 filtering and returned species level identification of *Corynebacterium*. Alignment and phylogenetic analysis identified four species, *C. accolens* (10/18), *C. mastitidis* (1/18), *C. tuberculostericum* (5/18) and *C. simulans* (2/18). The small sample size and skewed case-control status limited any meaningful tests for association.

### Relationship between ocular microbial community and host genotype

We have previously found an association between HLA-C2 copy number, *KIR2DL2*/*KIR2DL3* heterozygosity and conjunctival scarring.^7^ *KLRC2* (NKG2C) was also investigated as it is an activating receptor of NK cells and T cells that could potentially impact bacterial community structure or host responses. Polymorphisms in expression of this receptor have previously been shown not to be associated with trachoma however.^42^ Relationships between these host genetic factors and ocular microbiome in adults were determined by Hill numbers and nonmetric-multidimensional scaling (nMDS) of complete microbial communities. No significant associations were identified (Table 3). Details of host genotype by trachomatous disease stage are available in the supplement (Supplementary Table 6).

**Table 3.**
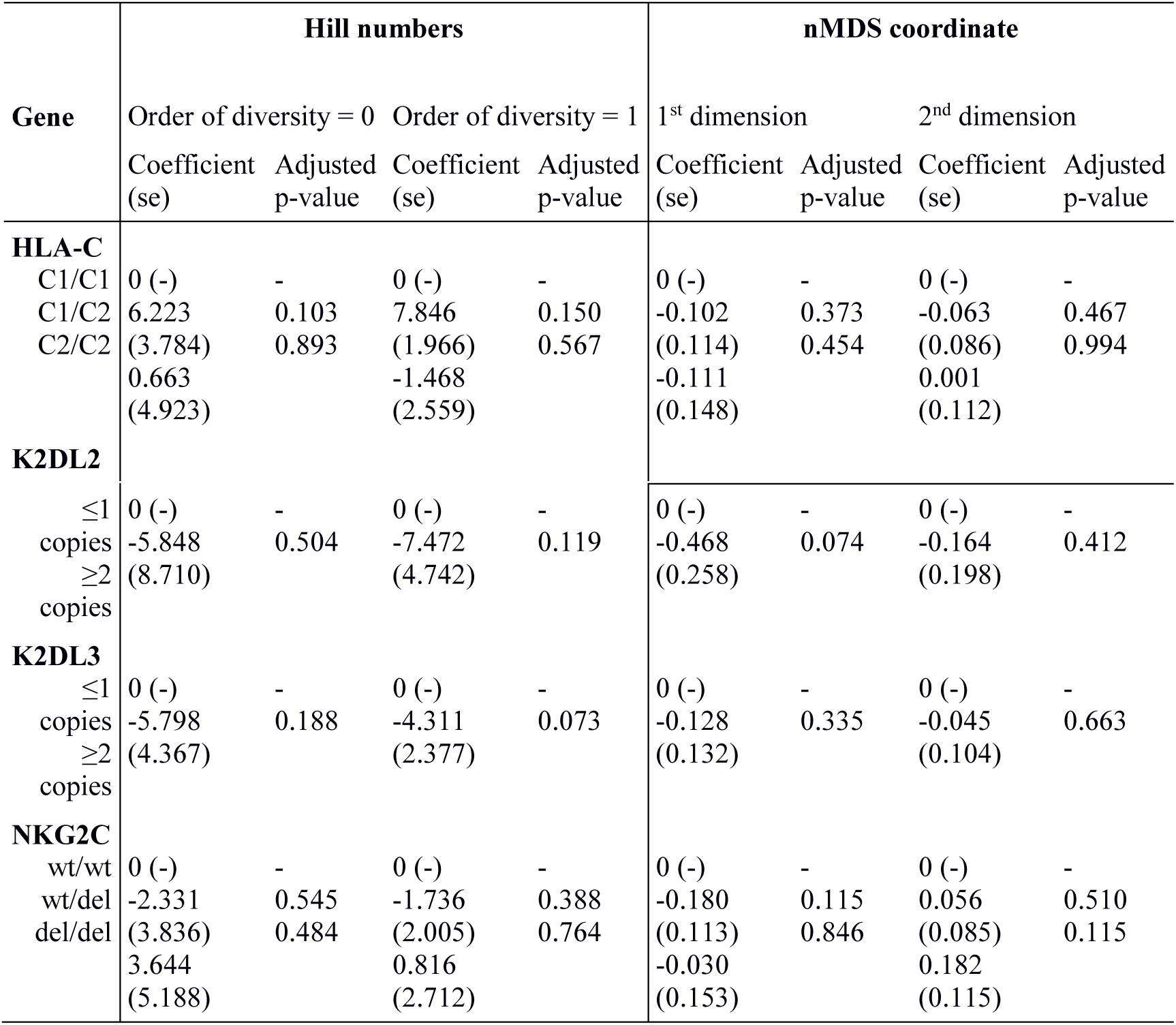
Associations between ocular microbiome and host genotype

### Relationship between gene expression, microbial community and evidence of trachomatous disease

Previous studies have implicated interplay between host gene expression and microbiome in disease pathogenesis^33-36^. We now evaluate the interplay of host ocular microbiome and conjunctival gene expression using a linear regression of GE on relative abundance, with previously detailed adjustments. In children, univariate analyses identified a number of genes associated with abundance of *Haemophilus* (Table 4). *S100A7, SRGN* and *TLR4* were upregulated in children with increased abundance of *Haemophilus*. These genes were all within module ATEM2, the expression of which was also significantly upregulated in children with increased abundance of *Haemophilus* (adj.p = 0.0001, coef = 7.777 [se = 1.898]). Visualisation of the relationship between gene expression and microbiome suggested the association between *Haemophilus* abundance and expression levels of ATEM2 was not linked to active trachoma (Figure 6). No further associations between microbiome and GE were identified.

**Table 4.** Significant associations between host gene expression and ocular microbes in children

| Gene | Genera | Adjusted p-value | Coefficient (se) |
| --- | --- | --- | --- |
| S100A7 | *Haemophilus* | 1.5x10<sup>-5</sup> | 7.15 (1.52) |
| SRGN | *Haemophilus* | 3.4x10<sup>-5</sup> | 3.04 (0.68) |
| TLR4 | *Haemophilus* | 6.2x10<sup>-5</sup> | 2.43 (0.57) |
| miR-155 | *Haemophilus* | 0.00016 | -6.10 (1.51) |

**Figure 6.**
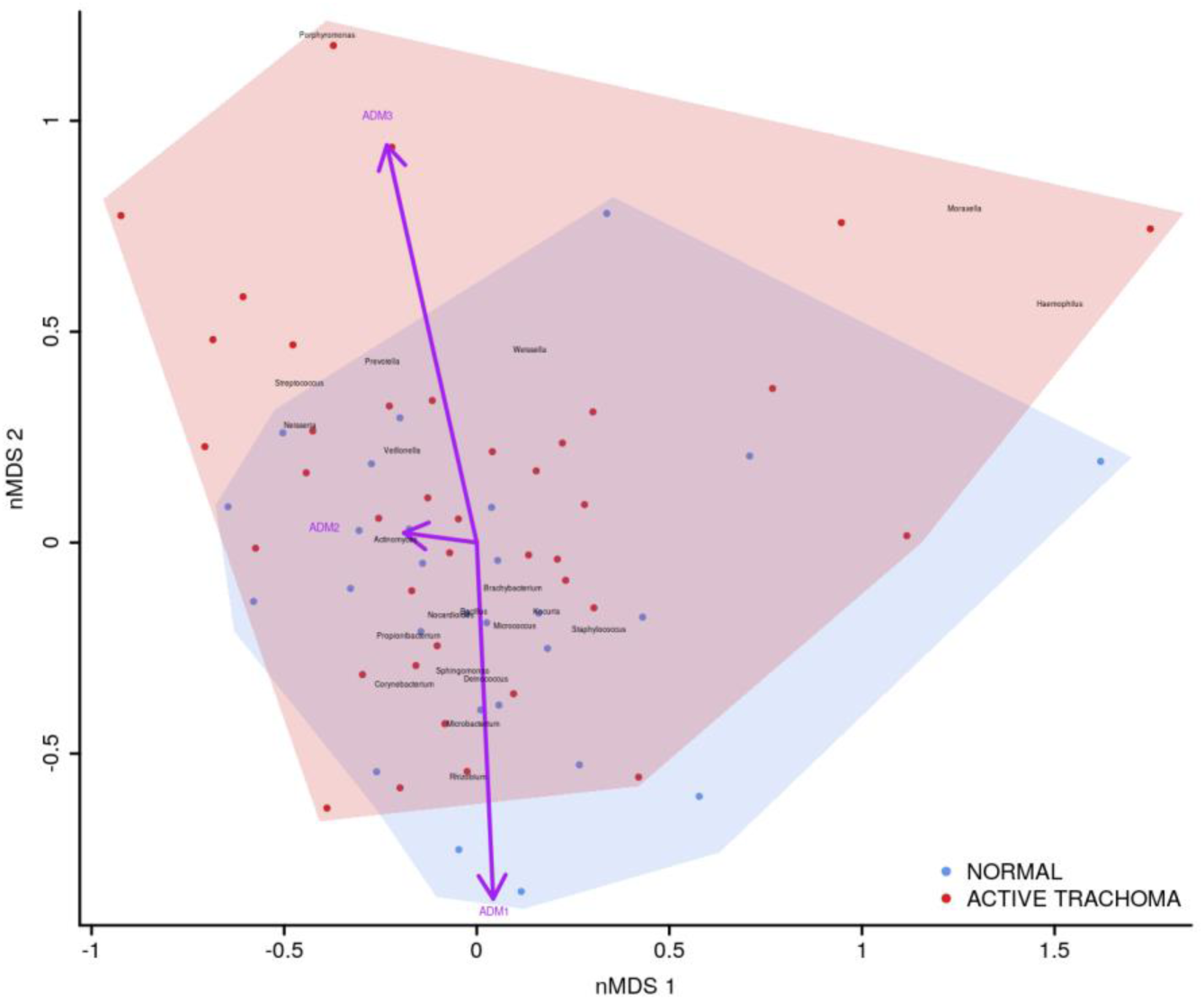
Plot of the ocular microbiome, modular conjunctival gene expression and trachomatous disease in children. Nonmetric multidimensional scaling of the complete ocular microbial community was used to position samples (points), enrichment of genera with relative abundance >1% are shown (black text). Arrows represent conjunctival gene expression modules (purple text), arrow coordinates indicate increased expression in surrounding samples. Shaded areas highlight the distribution of active trachoma (red) and healthy control (blue) samples.

In adults, univariate analyses identified 3 genes upregulated with increasing abundance of *Corynebacterium* (Table 5). These genes were all within module STEM4, expression of which was also significantly upregulated with increased abundance of *Corynebacterium* (adj.p = 0.0003, coef = 2.472 [se = 0.672]). Visualisation of the relationship between gene expression and microbiome showed greater separation of scarring trachoma and healthy individuals than observed in children with active trachoma and healthy individuals (Figures 6 & 7). Adults with ST clustered in the negative space of the first dimension of nMDS. Expression of STEM4 and abundance of *Corynebacterium* were both increased in this space populated exclusively by individuals with ST, suggesting the association between them is linked to ST. No further associations between microbiome and GE were identified.

**Table 5.** Significant associations between host gene expression and ocular microbes in adults

| Gene | Genera | Adjusted p-value | Coefficient (se) |
| --- | --- | --- | --- |
| CEACAM1 | *Corynebacterium* | 0.0009 | 1.37 (0.39) |
| MUC1 | *Corynebacterium* | 0.0001 | 1.25 (0.32) |
| MUC4 | *Corynebacterium* | 0.0003 | 1.48 (0.39) |

**Figure 7.**
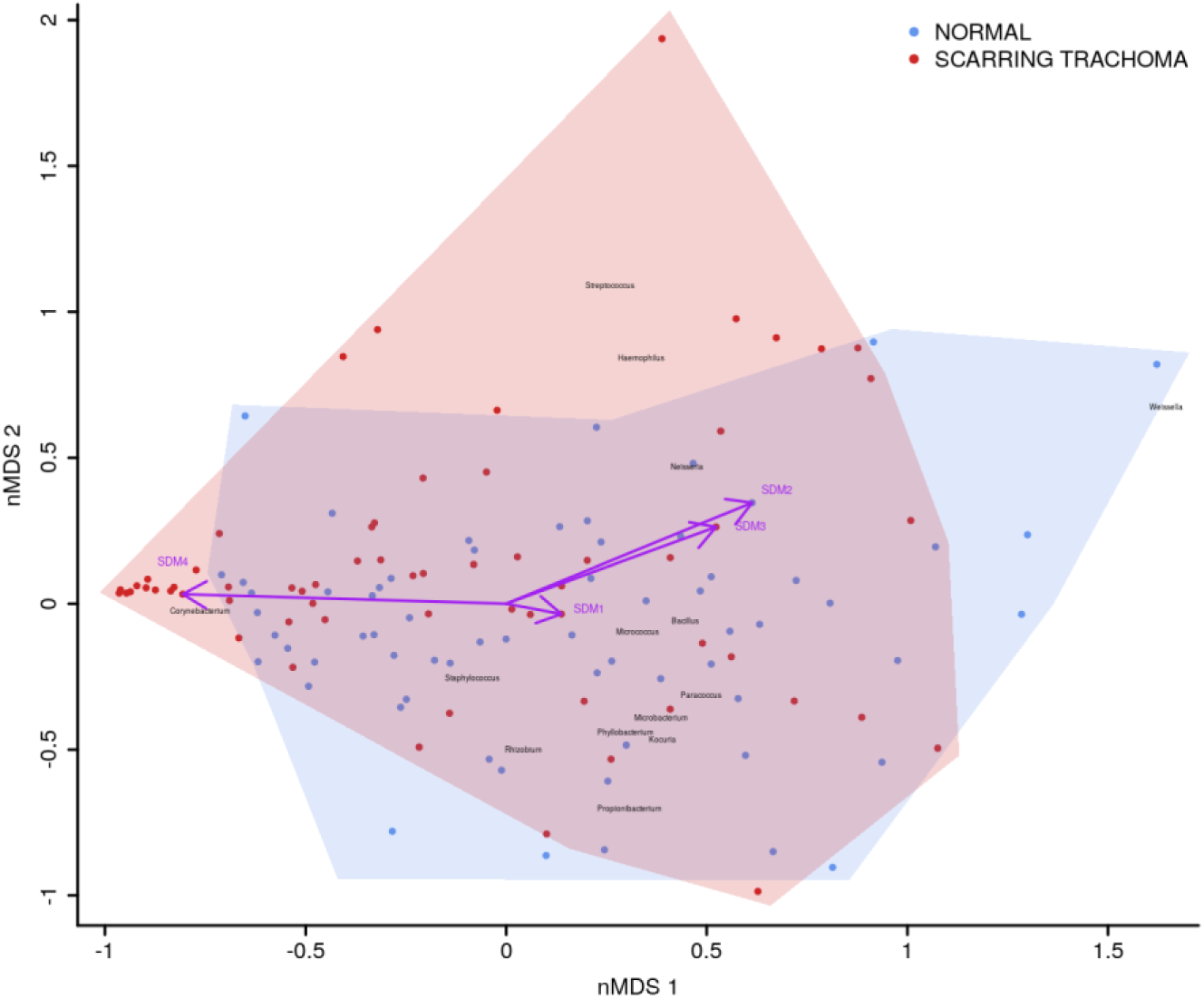
Plot of the ocular microbiome, modular conjunctival gene expression and trachomatous disease in adults. Nonmetric multidimensional scaling of the complete ocular microbial community was used to position samples (points), enrichment of genera with relative abundance >1% are shown (black text). Arrows represent conjunctival gene expression modules (purple text), arrow coordinates indicate increased expression in surrounding samples. Shaded areas highlight the distribution of scarring trachoma (red) and healthy control (blue) samples.

## Discussion

This is the first study that has examined conjunctival gene expression in the context of a culture-free characterization of the ocular microbiome in active and scarring trachoma in comparison with healthy controls. We found pro-inflammatory, antimicrobial and tissue-remodeling gene expression was associated with AT and ST, as previously described.^5, 6, 13^ Novel comparisons between AT and ST highlighted the importance of pro-inflammatory responses and tissue-remodeling in these pathogeneses. ST was associated with reduced microbial diversity and dominance of *Corynebacterium* whilst AT was not associated with detectable changes in the microbiota, both of which have been shown previously.^9-11^ Disease-associated changes in gene expression were strongly associated with imbalances in the microbiota, notably with increased abundance of *Haemophilus* in children and *Corynebacterium* in adults. In children, this relationship was independent of AT. Conversely, in adults, the relationship between changes in gene expression and *Corynebacterium* abundance were strongly associated with ST. Host genotype of targeted innate immune genes were not associated with ocular microbiome diversity or composition.

We found patterns of differential gene expression typical of AT and ST, as described previously in The Gambia and independently in other populations including Ethiopia and Tanzania^5, 6, 13^. In AT, we found anti-microbial peptides (S*100A7* and *DEFB*) had the largest significant up-regulation. *SPARCL1*, an inhibitor of extracellular matrix remodeling, was the most reduced transcript in both AT and ST, possibly indicating cellular remodeling of the conjunctiva taking place in both stages of disease. Upregulation of *IL17A/IL23A* and suppression of *MUC16* was also clearly evident in AT. This expression pattern, indicative of a type 17 response, is generally considered to be triggered by extracellular bacterial pathogens, fungi and some commensal bacterial species that bind to epithelial cells. These type 17-associated cytokines can be both beneficial (promoting mucosal barrier function) or detrimental (being associated with chronic inflammatory disorders). The gene expression pattern reflected in ST is also dominated by up-regulation of *DEFB* expression, accompanied by *MUC1/MUC4* and smaller differences in *IL23A* and *S100A7*, with little change in *IL17A* expression. Perhaps the most interesting observation is the reduction in *IL22* expression and the lack of *IL17A* differential expression in adults, expression of which are required for maintaining epithelial cell barrier health and recruitment of neutrophils, respectively. In ST, the epithelial cell layer is typically thinned and denuded in parts^49^ which is thought to be permissive or conducive to further bacterial colonisation and changes to the bacterial microbiome (dysbiosis).

Profiling both children with AT and adults with ST permits us to characterize the primary features of gene expression between the early and later stages of clinical disease. The largest differences between disease stages were the relatively reduced levels of expression of *IL22, IFNG* and miR-1285 in adults with ST compared to children with AT. This is consistent with a damaged epithelial cell layer, cellular proliferation and tissue re-organisation in the conjunctivae of adults. The relative absence of *IFNG* is consistent with previous transcriptome and targeted gene expression results in Ethiopian and Tanzanian adult trachoma cases with scarring or trichiasis.^5, 6, 13^ This lack of immune interferon suggests a lack of a dominant Th1 response in ST, unlike in AT, in which Th1 and type-1 interferon response profiles have been described as critical for clearance of *Ct* infection.^3, 50^

We found no evidence of significant changes in the ocular bacterial communities of children with AT compared to matched, healthy controls. Overall the prevalence of ocular *C. trachomatis* infection was low in AT cases, which is consistent with the observed decline in AT and infection prevalence seen in The Gambia in these districts prior to the mass distribution of Zithromax.^51^ The V1-V3 16S rRNA sequencing method used in this and our previous studies is not capable of resolving *Chlamydiae* taxa, which is a limitation of this study, however in AT cases there was no evidence of an overall dysbiosis.

The most substantial changes in relative abundance across each disease phenotype are in *Corynebacterium*. In children with AT there was a non-significant decrease in relative abundance whilst in adults with ST there was a significant increase. In adults with ST, there is almost a 1/3 increase in *Corynebacterium* relative abundance compared to controls. This is consistent with culture results in trachoma where adults with ST are consistently found to have an increased proportion of *Corynebacterium* isolated.^9^ A longitudinal study of progressive scarring disease suggested that presence of commensal bacteria by microbiological culture, including *Corynebacterium* spp, is marginally associated with further development of disease, although the effect was enhanced when in the presence of an ocular pathogen.^11^

There are > 120 species within the *Corynebacterium* genus. This genus is almost always isolated in ocular culture and identified in sequence studies of both healthy and diseased conjunctiva. *Corynebacterium* is largely disregarded as either a commensal, skin contaminant, or a secondary infection without a significant role in disease. Speciation of *Corynebacterium* in healthy eyes is not frequently investigated; one of the few reported studies found 23 of 92 healthy individuals positive for *Corynebacterium*, consisting of 5 lipophilic species (*C. macginleyi*, C. *afermentans subsp. lipophilum, C. accolens*, unspeciated lipophilic corynebacteria and *C. jeikeium*).^52^ Our samples were dominated by *C. accolens* and *C. tuberculostericum*. The identification of *Corynebacterium* species not commonly resident on skin suggests that these are not contaminants from surrounding facial skin, but species which find a niche on the conjunctiva.

We found both individual genes and modules (combinations of genes) that were differentially expressed and associated with the relative abundance of a number of genera. In children, three transcripts (*S100A7, SRGN* and *TLR4*) and miR-155 expression were associated with presence of *Haemophilus*. These genes were members of ATEM2, an expression module with 24 genes; the expression of which is characteristic of innate responses to the microbiota. Combined analysis of gene expression, ocular microbiome, and disease status suggested that the association between *Haemophilus* abundance and ATEM2 expression was not related to AT. This supports the lack of independent association between *Haemophilus* abundance and AT. Increased ATEM2 expression in AT may be driven by a recently cleared bacterial infection or an unknown non-bacterial pathogen. It is also possible that previous infections have caused epigenetic changes which predispose individuals to inflammatory responses in the absence of classical stimuli. A larger sample size is required to investigate these hypotheses.

In adults, increased relative abundance of *Corynebacterium* was associated with increased expression of mucins (*MUC1/MUC4*) and the adhesion molecule *CEACAM1. CEACAM1* in particular is exploited in mucosal colonisation by both pathogenic and nonpathogenic bacteria, increased expression may support ocular expansion of otherwise transient bacteria.^53^ Numerous splice variants of both mucins exist.^54, 55^ The frequency of two *MUC1* variants have been implicated in dry eye disease through modulation of the local inflammatory response,^54^ it is possible changes in the frequency of variants may increase susceptibility to ST. These 3 genes are members of a larger module, STEM4, containing 12 genes whose expression is driven by the ocular microbiota. In this case, the majority of increased STEM4 expression was associated with increased relative abundance of *Corynebacterium*, which was further enhanced in ST. A proposed hypothesis from work in mice is that *C. mastiditis* is required to stimulate innate resistance^24^, whilst in humans *C. accolens* is able to competitively generate metabolites that inhibit the growth of ocular pathogens such as *S. pneumoniae*.^*25*^ In contrast, our data suggest *Corynebacterium* spp found in the ocular niche may contribute to altered mucin expression, along with bacterial and epithelial cell adhesion which are important factors in ST. To determine if increased *Corynebacterium* abundance is a promoter of or a result of ST requires longitudinal investigation.

The combined examination of the ocular microbiome and conjunctival host response suggests contrasting profiles in AT and ST. In children, in the absence of current *C. trachomatis* infection, ocular pathogens such as *Haemophilus* are prevalent and associate with damaging inflammatory responses that impair epithelial cell health. However, these pathogens are not independently associated with AT, suggesting there may be undiscovered factors promoting inflammation. In adults, expansion of a non-pathogenic or commensal bacterium such as *Corynebacterium*, at the cost of bacterial community diversity, is associated with innate responses thought to drive ST. This response in adults was associated with increased mucin expression and enhanced matrix adhesion of epithelial cells. Enhanced cell matrix adhesion could contribute to fibrosis, whilst increased mucin expression may modulate inflammatory responses. Longitudinal studies are critical to the further understanding of progression of AT to ST. In AT, longitudinal studies are needed to understand how long inflammatory responses are sustained after clearance of an infection. In ST, longitudinal studies are required to investigate innate responses and *Corynebacterium* abundance and whether they are drivers or outcomes of scarring.

## Data sharing

Anonymised 16S sequencing data are available from the Sequence Read Archive (SRA) at the National Center for Biotechnology Information (NCBI) under accession numbers PRJNA248889 and PRNA515408.

## Acknowledgements

We express our thanks to the study participants, field team and support services in The Gambia. We also extend our thanks to Dr David Nelson and Dr Evelyn Toh of Indiana University for providing 16S rRNA primer sequences and valuable advice.

## Author contributions

Study design: DCWM, RLB, MJB, ChR, SEB, MJH

Sample collection: PM, HJ, TD, SEB, MJH

Data collection: HP, CDP, JH, TD, AG

Data analysis: HP, CDP, JH, TD, AG, ChR, SEB, MJH

Manuscript preparation: HP, CDP, JH, PM, HJ, TD, AG, DCWM, RLB, MJB, ChR, SEB, MJH

## Contributions to the field

Ocular *Chlamydia trachomatis* infection causes trachoma, the leading infectious cause of blindness worldwide. Previous studies have identified local inflammatory immune responses and associated changes in the bacterial flora of the eye as important in the disease process. Additionally, studies from other diseases have suggested certain constituents of the bacterial flora may protect against pathogenic infections. This study investigates the impact of local immune responses and changes in the bacterial flora on early and later stages of trachoma. We found evidence of strong inflammatory responses characteristic of active trachoma in early disease, with later stages dominated by damaging changes in local tissue health indicative of scarring disease. Coincidence of inflammatory immune responses and expansion of *Corynebacterium* in the later stages of trachoma, suggests a pathogenic role for non-chlamydial bacteria such as *Corynebacterium*. This study found that inflammatory responses are strongly associated with trachomatous inflammation and scarring. The association of specific non-chlamydial bacteria with damaging immune responses and disease supports further of their role in trachoma pathogenesis. The common theme of suppression of tissue re-epithelialisation in early and late stages of trachoma suggests that stimulating immune responses that promote tissue homeostasis and resolution of inflammation will be important in limiting scarring damage in trachoma and other inflammatory conjunctival diseases.

