## Supplementary Information for "Conjunctival microbiome-host responses are associated with impaired epithelial cell health in both early and late stages of trachoma"

### Supplementary Tables

Supplementary Table 1. TLDA card details

| General category* | Target |
|---|---|
| Anti-microbial peptides[1-4] | DEFB4B_DEFB4A, S100A7, |
| Cell adhesion/growth regulation[1] | CEACAM1, CD53 |
| Cell cycle/proliferation[1] | CDK13, PRDX3, TP53 |
| Cell markers[1] | CD247, KLRF1 |
| Cytokines/chemokines[1, 2, 4] | CCL2, IFNG, IL12B, IL17A, IL22, IL23A, TNFSF15 |
| Mucins/matrix modifiers[2, 4, 5] | MMP7, MUC1, MUC4, MUC5AC, MUC7, MUC16, SPARCL1 |
| Pattern recognition/inflammasome[1] | NLRP1, NLRP3, NLRP6, NOD1, NOD2, RIPK2, TLR4, TLR6 |
| Regulators/signaling pathways[1] | CHD8, COMMD6, MAPK14, MYD88, NFKB1, PDL1, SRGN, TRAF6 |
| Regulatory elements[6, 7] | miR-147B, miR-155, miR-184, miR-1285 |
| Response to microbiota[1, 4] | ALOX5AP, ANAX1, BCL2, CD40, REL, TNFRSF1A, TNFRSF1B |

**Reference for evidence of differential regulation in trachoma*

Supplementary Table 2. Gene expression sample demographics

| Phenotype | Children (< 16 years of age) | | | Adults (>= 16 years of age) | | |
|---|---|---|---|---|---|---|
| | Healthy n = 32 (41%) | Active trachoma n = 46 (59%) | Adjusted p-value | Healthy n = 69 (47%) | Scarring trachoma n = 78 (53%) | Adjusted p-value |
| **Male (%)** | 18 (56) | 29 (39) | 0.713 | 22 (32) | 16 (21) | 0.167 |
| **Median age (range)** | 6 (1-13) | 5 (1-13) | 0.291 | 47 (16-87) | 50 (16-84) | 0.063 |
| **Sample collection [n(%]** | | | 0.884 | | | 1.000 |
| dry season | 5 (16) | 9 (20) | | 51 (74) | 57 (73) | |
| wet season | 27 (84) | 37 (80) | | 18 (26) | 21 (27) | |
| **Ethnicity [n(%)]** | | | 0.439 | | | 0.495 |
| Jola | 5 (16) | 6 (13) | | 23 (33) | 23 (29) | |
| Mandinka | 14 (44) | 17 (37) | | 30 (43) | 43 (55) | |
| Wolof | 10 (31) | 12 (26) | | 6 (9) | 4 (5) | |
| Other | 3 (9) | 11 (24) | | 10 (14) | 8 (10) | |
| **Administrative district [n(%)]** | | | 0.170 | | | 0.456 |
| Basse | 0 (0) | 0 (0) | | 7 (10) | 9 (12) | |
| Brikama | 10 (31) | 15 (33) | | 34 (49) | 39 (50) | |
| Janjanbureh | 1 (3) | 0 (0) | | 1 (1) | 0 (0) | |
| Kanifing | 0 (0) | 0 (0) | | 1 (1) | 1 (1) | |
| Kerewan | 0 (0) | 3 (7) | | 6 (9) | 4 (5) | |
| Kuntaut | 2 (6) | 0 (0) | | 3 (4) | 4 (5) | |
| Mansa Konko | 19 (59) | 28 (61) | | 8 (12) | 21 (27) | |
| Unknown | 0 (0) | 0 (0) | | 9 (13) | 0 (0) | |

Supplementary Table 3. Univariate and modular gene expression in children

| **Module** *putative function* | **Target** | **Fold-change*** | **Adjusted p-value** |
|---|---|---|---|
| ATEM1** *cellular adhesion and division* | ANXA1 | 1.148 | 0.636 |
| | BCL2 | 1.146 | 0.689 |
| | CD247 | 1.212 | 0.520 |
| | CEACAM1 | 1.326 | 0.810 |
| | IFNG | 1.174 | 0.260 |
| | IL12B | 0.507 | 0.104 |
| | IL22 | 0.498 | 0.004 |
| | KLRF1 | 1.930 | 0.355 |
| | MMP7 | 1.090 | 0.881 |

| | | | |
|---|---|---|---|
| | miR-1285 | 0.494 | 0.372 |
| | miR-147B | 1.010 | 0.881 |
| | miR-155 | 1.300 | 0.101 |
| | miR-184 | 0.934 | 0.668 |
| ATEM2<br>*innate response to microbiota* | ALOX5AP | 0.871 | 0.288 |
| | CCL2 | 0.901 | 0.347 |
| | CD274 | 0.971 | 0.756 |
| | CD53 | 0.664 | 0.017 |
| | COMMD6 | 0.559 | 0.101 |
| | DEFB4B_DEFB4A | 1.122 | 0.347 |
| | IL17A | 0.890 | 0.421 |
| | IL23A | 1.349 | 0.101 |
| | MUC7 | 0.952 | 0.756 |
| | NLRP3 | 0.797 | 0.143 |
| | NLRP6 | 1.066 | 0.689 |
| | NOD2 | 1.028 | 0.843 |
| | RIPK2 | 0.935 | 0.602 |
| | S100A7 | 0.907 | 0.602 |
| | SRGN | 0.709 | 0.003 |
| | TLR4 | 1.300 | 0.142 |
| | TLR6 | 1.849 | 0.118 |
| | TNFRSF1B | 1.520 | 0.003 |
| | TNFSF15 | 1.096 | 0.520 |
| ATEM3<br>*suppression of epithelial cell expansion/recovery* | CD40 | 1.132 | 0.756 |
| | CDK13 | 1.290 | 0.243 |
| | CHD8 | 1.602 | 0.041 |
| | MAPK14 | 1.395 | 0.101 |
| | MUC1 | 1.416 | 0.055 |
| | MUC16 | 1.323 | 0.229 |
| | MUC4 | 0.950 | 0.881 |
| | MUC5AC | 1.021 | 0.881 |
| | MYD88 | 2.077 | 0.003 |
| | NFKB1 | 1.139 | 0.494 |
| | NLRP1 | 2.921 | 0.024 |
| | NOD1 | 2.587 | 0.003 |
| | PRDX3 | 2.199 | 0.006 |
| | REL | 1.569 | 0.143 |
| | SPARCL1 | 1.047 | 0.750 |
| | TNFRSF1A | 1.324 | 0.103 |
| | TP53 | 3.919 | 0.005 |
| | TRAF6 | 0.198 | 0.003 |

**Fold-change in expression from healthy controls to active trachoma cases*
***ATEM – active trachoma expression module*

### Supplementary Table 4. Univariate and modular gene expression in adults

| **Module**<br>*putative function* | **Target** | **Fold-change*** | **Adjusted p-value** |
|---|---|---|---|
| STEM1**<br>*epithelial health* | COMMD6 | 1.148 | 0.342 |
| | IFNG | 1.146 | 0.010 |
| | IL12B | 1.212 | 0.495 |
| | IL22 | 1.326 | 0.006 |
| | KLRF1 | 1.174 | 0.668 |
| | MUC7 | 0.507 | 0.077 |
| | TNFSF15 | 0.498 | 0.008 |
| | miR-147B | 1.930 | 0.550 |
| | miR-155 | 1.090 | 0.275 |
| | miR-184 | 0.494 | 0.902 |
| STEM2<br>transcriptional control of innate sensing | BCL2 | 1.010 | 0.687 |
| | CD247 | 1.300 | 0.146 |
| | CD40 | 0.934 | 0.061 |
| | CDK13 | 0.871 | 0.797 |
| | CHD8 | 0.901 | 0.004 |

| | | | |
|---|---|---|---|
| | MAPK14 | 0.971 | 0.495 |
| | MUC16 | 0.664 | 0.535 |
| | MUC5AC | 0.559 | 0.294 |
| | MYD88 | 1.122 | 0.036 |
| | NLRP1 | 0.890 | 0.832 |
| | NLRP3 | 1.349 | 0.391 |
| | NOD1 | 0.952 | 0.239 |
| | PRDX3 | 0.797 | 0.130 |
| | REL | 1.066 | 0.130 |
| | RIPK2 | 1.028 | 0.294 |
| | TNFRSF1A | 0.935 | 0.888 |
| | TP53 | 0.907 | 0.271 |
| | TRAF6 | 0.709 | 0.169 |
| STEM3 *cellular proliferation and activation* | ALOX5AP | 1.300 | 0.130 |
| | CCL2 | 1.849 | 0.787 |
| | CD53 | 1.520 | 0.172 |
| | NFKB1 | 1.096 | 0.019 |
| | NLRP6 | 1.132 | 0.294 |
| | NOD2 | 1.290 | 0.106 |
| | SRGN | 1.602 | 0.124 |
| | TLR4 | 1.395 | 0.259 |
| | TLR6 | 1.416 | 0.888 |
| | TNFRSF1B | 1.323 | 0.943 |
| | miR-1285 | 0.950 | 0.007 |
| STEM4 *response to microbiota* | ANXA1 | 1.021 | 0.124 |
| | CD274 | 2.077 | 0.275 |
| | CEACAM1 | 1.139 | 0.137x10$^{6}$ |
| | DEFB4B_DEFB4A | 2.921 | 0.003 |
| | IL17A | 2.587 | 0.457 |
| | IL23A | 2.199 | 0.004 |
| | MMP7 | 1.569 | 0.004 |
| | MUC1 | 1.047 | 0.110x10$^{5}$ |
| | MUC4 | 1.324 | 0.137x10$^{7}$ |
| | S100A7 | 3.919 | 0.463 |
| | SPARCL1 | 0.198 | 0.001 |

**Fold-change in expression from healthy controls to active trachoma cases*
***STEM – scarring trachoma expression module*

### Supplementary Table 5. 16S sample demographics

| Phenotype | Children (< 16 years of age) | | | Adults (>= 16 years of age) | | |
|---|---|---|---|---|---|---|
| | Healthy (31[43%]) | Active trachoma (41[57%]) | Adjusted p-value | Healthy (105[45%]) | Scarring trachoma (130[55%]) | Adjusted p-value |
| **Male (%)** | 18 (58) | 26 (63) | 0.828 | 31 (24) | 34 (26) | 0.669 |
| **Median age (range)** | 5 (1-14) | 5 (1-12) | 0.382 | 52 (16-87) | 53 (16-84) | 0.188 |
| **Sample collection [n(%]** | | | 1.000 | | | 1.000 |
| dry season | 4 (13) | 6 (15) | | 78 (74) | 97 (75) | |
| wet season | 27 (87) | 35 (85) | | 27 (26) | 33 (25) | |

| | | | | | | |
|---|---|---|---|---|---|---|
| **Ethnicity [n(%)]** | | | 0.705 | | | 0.763 |
| Jola | 6 (19) | 6 (15) | | 36 (34) | 44 (34) | |
| Mandinka | 12 (39) | 13 (32) | | 43 (41) | 59 (45) | |
| Wolof | 8 (26) | 11 (27) | | 9 (9) | 7 (5) | |
| Other | 5 (16) | 11 (27) | | 17 (16) | 20 (15) | |
| **Administrative district [n(%)]** | | | 0.530 | | | 0.917 |
| Basse | 0 (0) | 0 (0) | | 8 (8) | 12 (9) | |
| Brikama | 11 (35) | 13 (32) | | 57 (54) | 71 (55) | |
| Janjanbureh | 0 (0) | 0 (0) | | 2 (2) | 3 (2) | |
| Kanifing | 0 (0) | 0 (0) | | 1 (1) | 1 (1) | |
| Kerewan | 0 (0) | 1 (2) | | 8 (8) | 10 (8) | |
| Kuntaut | 1 (3) | 0 (0) | | 5 (5) | 5 (4) | |
| Mansa Konko | 19 (61) | 27 (66) | | 14 (13) | 28 (22) | |
| Unknown | 0 (0) | 0 (0) | | 10 (9) | 0 (0) | |

### Supplementary Table 6. Host genotype in adults by case-control status

| | **Adults (>= 16 years of age)** | | |
|---|---|---|---|
| **Phenotype** | Healthy (121[43%]) | Scarring trachoma (158[57%]) | Adjusted p-value |
| **KIR2DL2 copy number [n(%)]** | | | 0.412 |
| 0 | 45 (37) | 71 (45) | |
| 1 | 33 (27) | 45 (28) | |
| 2 | 6 (5) | 4 (3) | |
| Unknown | 37 (31) | 38 (24) | |
| **KIR2DL3 copy number [n(%)]** | | | 0.492 |
| 0 | 14 (12) | 14 (9) | |
| 1 | 47 (39) | 77 (49) | |
| 2 | 22 (18) | 31 (20) | |
| Unknown | 38 (31) | 36 (23) | |
| **HLA-C genotype [n(%)]** | | | 0.228 |
| C1 | 27 (22) | 47 (30) | |
| C1C2 | 50 (41) | 64 (41) | |
| C2 | 26 (22) | 24 (15) | |
| Unknown | 18 (15) | 23 (15) | |
| **NKG2C genotype [n(%)]** | | | 0.605 |
| WT/WT | 47 (39) | 53 (34) | |
| WT/DEL | 55 (45) | 77 (49) | |
| DEL/DEL | 16 (13) | 25 (16) | |
| Unknown | 3 (3) | 3 (2) | |

### Supplementary Figures

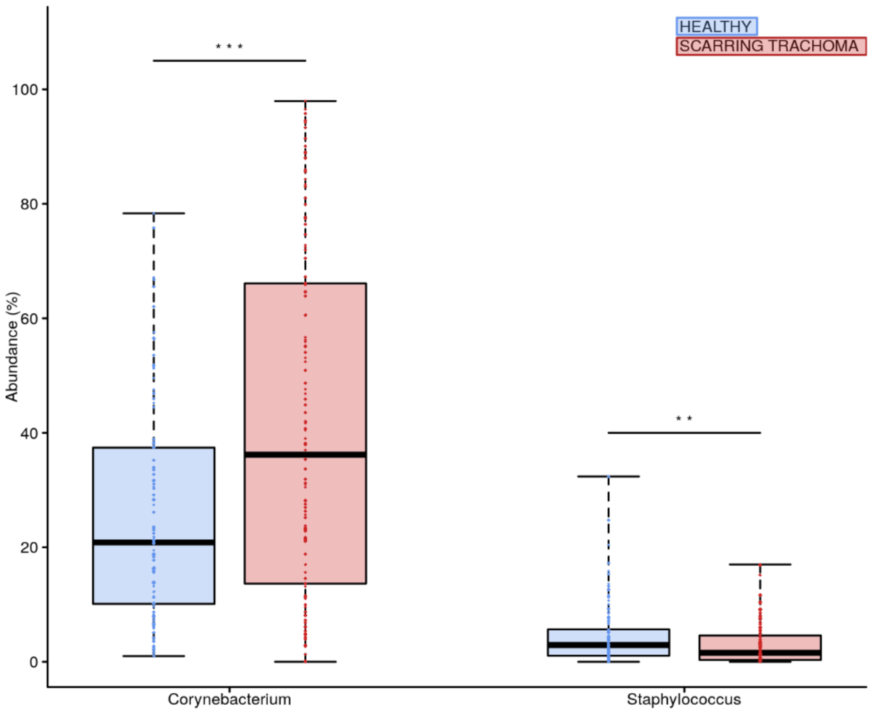


Supplementary Figure 1. Univariate analysis of relative abundance of genera by case-control status in adults.

Genera with significantly different relative abundance between scarring trachoma cases (red) and healthy controls (blue) are shown. Boxes represent the interquartile range, with median indicated (blue line). Outer bars represent the range. P-values were considered significant at <0.05 and are denominated as follows: * p<0.05; ** p<0.01; *** p<0.001.
